# High-Resolution Characterization of Human Brain Cortex with High-Fidelity Spatial Transcriptomic Slides (HiFi-Slides)

**DOI:** 10.1101/2023.06.12.544625

**Authors:** Tianyang Xu, Ekko Zhu, Chi Zhang, Riccardo Calandrelli, Pei Lin, Sheng Zhong

**Affiliations:** Shu Chien-Gene Lay Department of Bioengineering, Jacobs School of Engineering, University of California, San Diego, 9500 Gilman Drive, La Jolla

**Author notes:** Correspondence: Tianyang Xu, Sheng Zhong.

**Keywords:** Spatial Transcriptomic, Alzheimer’s disease, High Resolution, Next Generation Sequencing

## Abstract

Spatial transcriptomic tools and platforms help researchers to inspect tissues and cells with fine details of how they differentiate in expressions and how they orient themselves. With the higher resolution we get and higher throughput of expression targets, spatial analysis can truly become the core player for cell clustering, migration study, and, eventually, the novel model for pathological study. We present the demonstration of HiFi-slide, a whole transcriptomic sequencing technique that recycles used sequenced-by-synthesis flow cell surfaces to a high-resolution spatial mapping tool that can be directly applied to tissue cell gradient analysis, gene expression analysis, cell proximity analysis, and other cellular-level spatial studies.

## Introduction

### HiFi-Slides Spatial Transcriptomic Applications

Developments and applications of spatial transcriptomic tools, such as spatial barcoded sequencing(1, 2) and in-situ sequencing(3–5) are powerful tools for studying tissue and disease pathology. The efforts to define the fine cellular level gene expressions convoluting with other molecular, pathological, and histochemical annotations or tissue maps advance the researchers’ ability to establish human organ references, disease pathways, and multi-omic data analysis (6). We have listed and compared the current popular techniques for spatial transcriptomic tools in Table 1. The High-Fidelity Spatial Transcriptomic Slides (HiFi-Slides) we demonstrated here were designed to address the need for an affordable solution for high-resolution total RNA spatial transcriptomic analysis.

During the development of this project, Cho et al. (5) reported the method of Seq-Scope with flow cells generated by a randomized library. Designed independently and different from the previous flow cell-based high-resolution spatial technique Seq-Scope, the HiFi-slide technique used a pre-qualified recycled flow cell instead of generating new flow cells with designed random libraries when both techniques utilize the high-density of unique barcodes found on sequencing-by-synthesize flow cells. Based on three core demands of new spatial transcriptomic tools (7): affordability, high throughput, and high resolution, the HiFi-slides technique turns recycled Illumina sequencing flow cells into spatial transcriptomic capture slides with terminal sequence modifications shown in Figure 1A and B, therefore, can have unique spatial barcodes that distributed within the distance within single cell level, available for gene expression mapping (Figure 1C).

**Fig 1.**
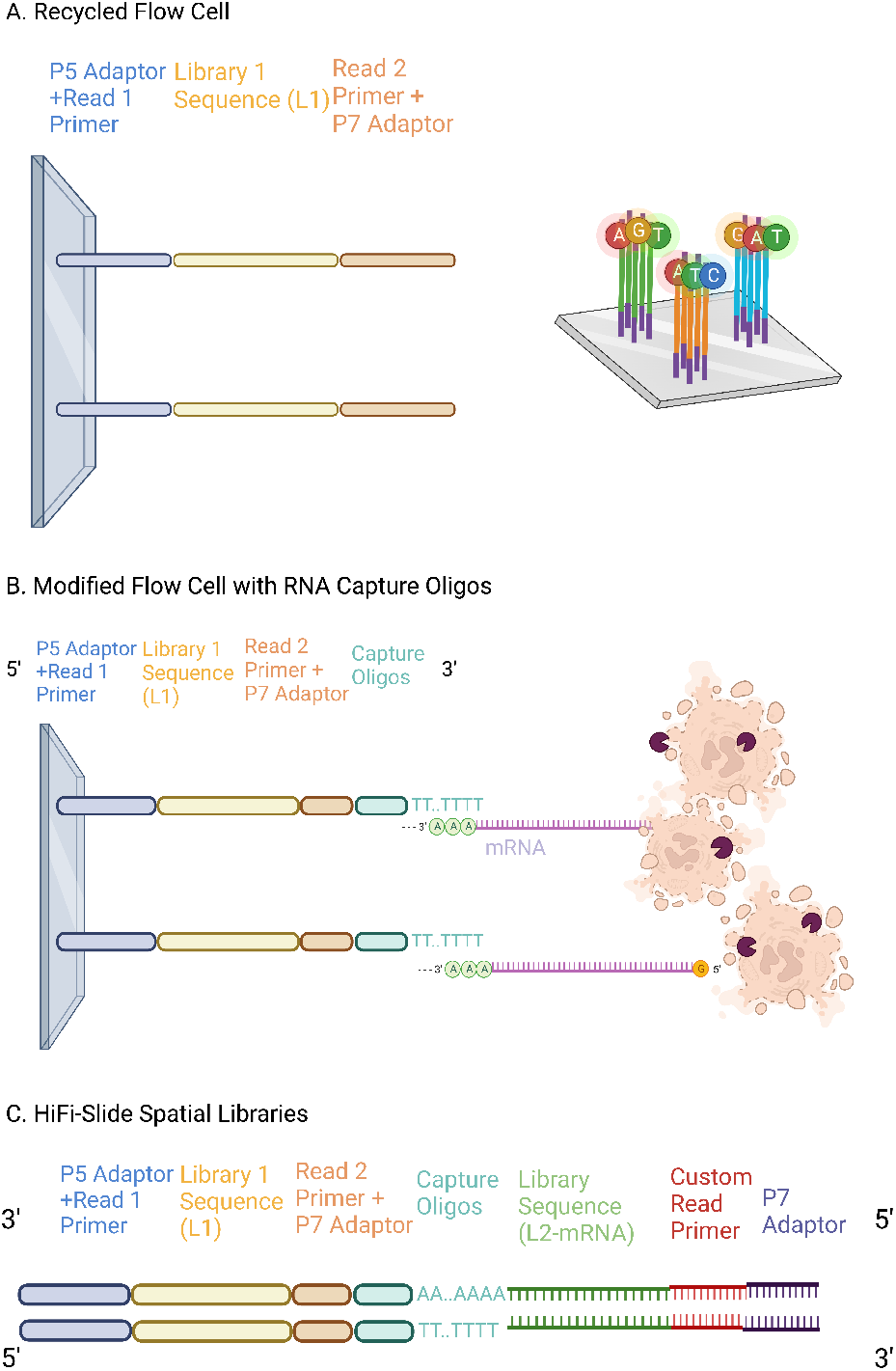
Demonstration of Sequences in HiFi-Slide. A: After the first round of sequencing, the strands synthesized on the flow cell are used as the barcodes with known coordination. B: Terminal modifications and ligation allow the strands on the slide to capture RNA released from the permeabilized cells and tissues. C: The complete library of HiFi-Slides has the P5 end of spatial barcodes and the P7 end of captured RNA fragments. A custom read primer 2 is required for the sequencing.

The sequencing-by-synthesis platform will generate millions of clusters on the surface of a sequencing flow cell (8). Each cluster has the same DNA sequences synthesized during the sequencing process, which will serve as the spatial barcodes in the HiFi-slide sequence. The distances between different clusters vary among types of flow cells from 0.4 microns to a few microns. The recycled flow cell reduces the cost of microarray acquisitions, with even higher resolution than the current commonly used technologies such as Visium (9) and Slide-Seq platforms (10).

### Spatial Studies on Alzheimer’s Disease (AD)

As one of the most prevalent forms of dementia, Alzheimer’s disease (AD) currently impacts millions of people world-wide. Due to its neurodegenerative nature, death typically occurs within 3 to 9 years after the initial diagnosis (11). Since the discovery of AD, continuous efforts have been invested in investigating its pathology and addressing the global financial and healthcare demands it imposes. Spatial transcriptomics, an innovative technique that combines spatial information with high-throughput gene expression analysis within the context of tissue architecture, has quickly been adopted in AD research to develop effective diagnostic and therapeutic strategies.

Previous spatial transcriptomic studies on AD using existing platforms have revealed valuable information, such as distinct gene expression patterns in specific regions of the brain associated with AD pathology, including amyloid beta metabolism, neuroinflammation, synaptic dysfunction, and neuronal loss (12). Transcriptomic profiles of different cell types, such as neurons, astrocytes, microglia, oligodendrocytes, and endothelial cells, have also been characterized to identify cell-type-specific gene expression alterations. The brain exhibits spatial heterogeneity, with certain regions, such as the cortex, being more susceptible to AD pathogenesis (12). Importantly, the findings from previous studies have been validated and integrated with other omics data, such as proteomics and genomics, revealing converging molecular pathways and mechanisms underlying AD. The spatial patterns of AD pathology change in the cell-type associated gene expressions, and the cell interactions in a certain region play key roles in the study of AD (13).

### Cell-type Specific Expression and Proximity Change in AD

AD pathologies, such as amyloid beta accumulation and neuroinflammation, induce transcriptional alterations within cell types and sub-types, which further leads to clinical symptoms (14). Reactive astrogliosis is the change of astrocytes, the hub of neuro-metabolism and homeostasis, in response to neurodegenerative diseases such as Alzheimer’s Disease or Parkinson’s Disease (15). In mouse models, astrocytes are reported to express more genes related to inflammation (16)and reduce the production of genes associated with neuronal support and protection(17). Other studies indicate that the number of oligodendrocytes and oligodendrocytes precursor cells is significantly decreased (18, 19), which impairs the myelin sheath regeneration and damage synaptic signaling (20). The recent development of single-cell sequencing techniques further discloses this cell heterogeneity (21). As the resident immune cells in the brain, microglia are activated in response to the neuroinflammation in neurodegenerative disease (22), which leads to a higher proportion of activated microglia (23) and a higher proportion of microglia subtype expressing inflammation-associated genes (24), but depletes the homeostatic microglia (25).

Though bulk analysis of single-cell RNA sequencing offered insights into cell type alteration and variation in AD, due to the spatial heterogeneity of AD pathology, such as amyloid-beta plaque, the neuronal cells also reflect the spatial pattern in the presence of AD pathology (26–28). Chen, et al. combined Spatial Transcriptomics and *in situ* Sequencing to reveal the increased co-expression pattern of astrocytes and microglia and the increase in oligodendrocytes as part of the amyloid-beta plaque environment (29). Chan, Chang, et al. utilized the 10X Visium slide to show an increased co-expression pattern within microglia cells (30).

However, few studies are yet reporting global cell proximity change in AD. Such discovery is limited to the study’s resolution and target throughput. Complex and heterogeneous tissues such as the brain cortex require spatial profiling will cellular resolution (micron level) and whole transcriptome coverage. The resolution of spatial transcriptomic techniques can impose certain constraints on the depth and specificity of analysis and eventually challenge the identification of subtle changes in gene expression patterns within specific cell types or the interaction and proximity shift between cells. The RNA target throughput decides the transcriptomic profiling depth, limiting the subsets of expression captured from the entire transcriptome (2), and ignoring cell type in low quantity and subtle changes in cell subtype variation, where important players implicated in AD pathology might remain undetected and underrepresented.

By applying the HiFi-slides technique to fresh frozen brain cortex samples of both none-AD (ND) and AD groups, we hope to use the ability of total RNA profiling and potentially submicron resolution mapping and advance the study of AD pathology in transcriptomic profiling depth, cell-type gene expression changes, and the detection sensitivity of the low-abundance transcripts.

## Results and Discussion

### Characteristics of HiFi-Slide Application

HiFi-slides were prepared from 6 used NextSeq 2000 P3 flow cells, with the read files but unknown sample information from the previous run. Each slide has 84 adjacent spatial tiles (6 columns and 14 rows) and 64 *mm*^2^ tissue application areas available. 12 fresh frozen slices from 6 donors were applied to HiFi-slides, including 1 AD patient sample and 5 ND control samples. Region of Interest (ROI) was selected from the tiles that were covered by the tissues.

The gene expression was then aligned and filtered by the tissue-covered region, as shown in Figure 2 in a 3 *mm*^2^ ROI region. 713,905 total spatially unique spots were mapped with gene-aligned expressions, and 160,369 spatial spots were visualized with a brain cortex cell-type associated gene filter, with 52,681 total genes expressed. We achieved an average of 23.8 unique addresses in 100 *um*^2^. For AD samples, 587,296 unique addresses with gene expressions were detected in the same area size, with 209,268 brain cell-type associated genes filtered and 19 unique addresses per 100 *um*^2^. HiFi-slide advanced in the gene target capture amount and spatial resolutions in the brain cortex application, approaching sub-micron resolution with a recycled flow cell.

**Fig 2.**
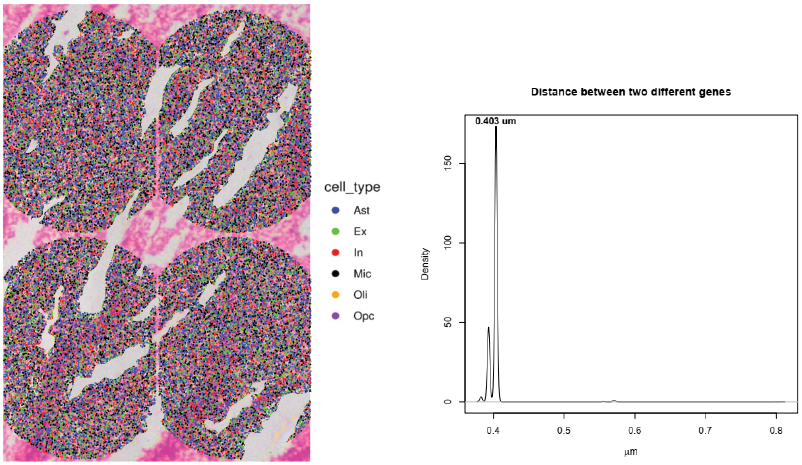
Left: Mapped gene expressions for four tiles (3 *mm*^2^) of ROI inside an ND sample. Colors of gene expression are associated with cell types shown in the legend. 713,905 unique spatial addresses were detected. 160,369 spatial spots contain brain cell-type-associated gene expressions. Right: Distribution of distances between two uniquely spatially resolved genes captured. Resolution range 0.382 to 0.807 microns with a mean distance of 0.402 microns

### Grey and White Matter Cell Gradient Analysis

Based on the high-resolution expressions captured with HiFi-slides, the cell-associated features of tissue can be inspected and visualized, such as the cell gradients. In an ND sample, 12 tiles (0.8 *mm*^2^ each tile)from the flow cell that contains the whole tissue were selected. A grey and white matter cell-type associated gene expression filter was applied. To discard genes expressed in a very low number of spots, we consider only those expressed in at least 1000 spots, giving a total of 10,855 genes for the section. To visualize the gradients, heatmaps of the number of spots expressing the selected gene set normalized by the total number of spots in each tile (Figure 3). The gradients of both white matter-associated cells and grey matter-associated cells confirm the gradient of the tissue cell orientations, which align with the tissue from the bottom left corner of the grey matter cells to the top right corner of the white matter cells.

**Fig 3.**
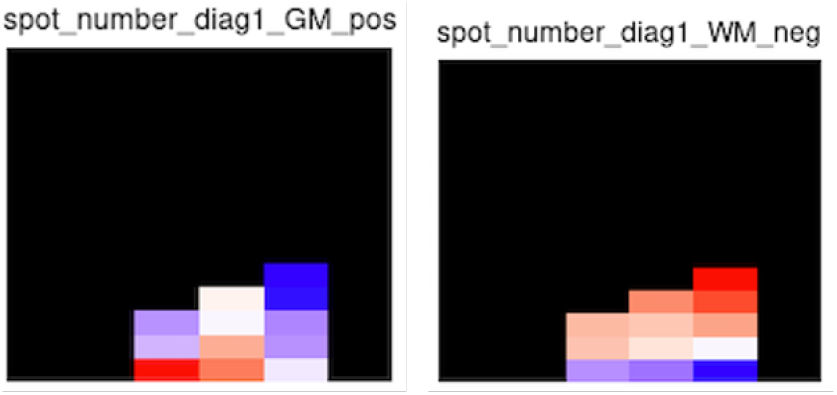
Gene expression gradient analysis shows the direction and the gradient change of the relative expression enrichment along the tissue direction; Gray matter (Left) in the bottom-left area, with strong depletion of gray matter on the top-right corner. The counterpart white matter cells (Right) show enrichment of white matter in the top-right corner but also with depletion at the bottom-left corner, which was enriched in grey matter. Bottom-right is shown as depleted as in grey matter.

### Cell-Type Associated Gene Expression Analysis

HiFi-slides can support the cellular level expression analysis to assist in the disease study. Cell-type associated expression filters and gene expression enrichment analysis was performed from the reads inside ROI tiles of both AD and ND samples (Figure. 4). The gene expression density in the spatial domain was analyzed and compared.

**Fig 4.**
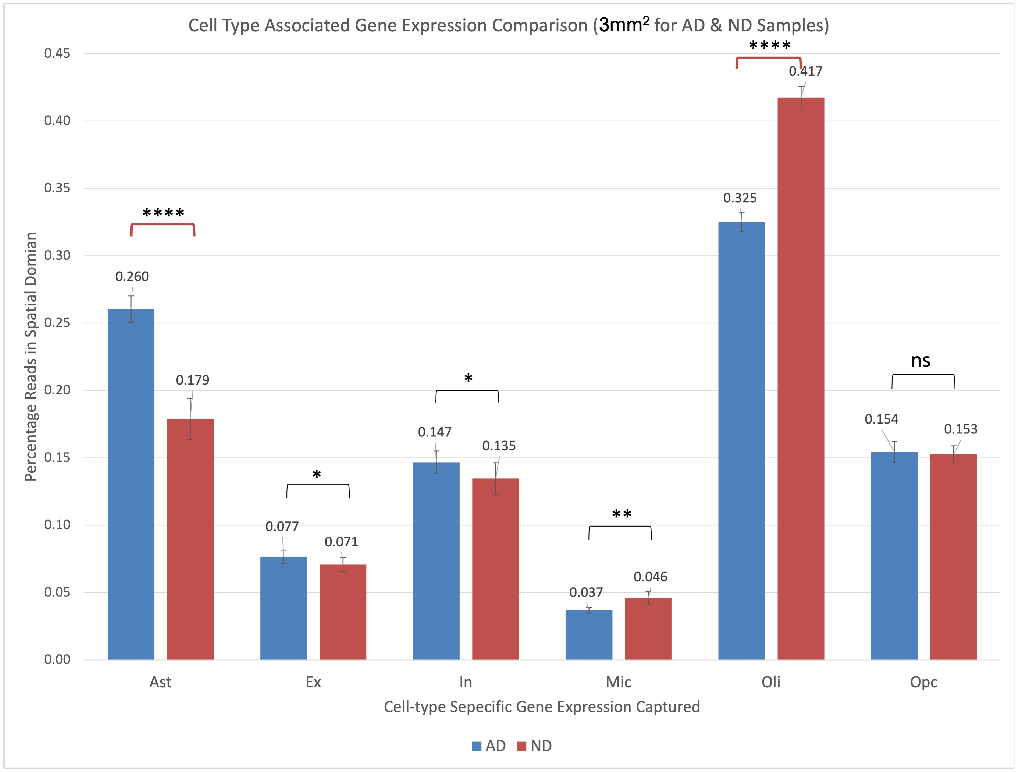
AD vs. ND cell-type associated gene expression differences on percentage expression in the spatial domain. Five standard deviations were shown

Astrogliosis results in the upregulation of astrocytes’ gene expression (31), which can be confirmed with HiFi-Slides analysis that the AD sample showed a 45% increase of astrocyte-associated gene expression in the spatial domain. As a cascading step for AD pathogenesis, such reactive astrogliosis may inhibit the growth of plaques as the neuroinflammatory response (32). Meanwhile, the gene expressions of Oligodendrocytes decreased by 22%, indicating disruption of its functions, as well as the myelin integrity, which always results in impairment of signal transmission between neurons, and the disruption of the supplement to axons (20). HiFi-slide results align with the observations with the spatial modality of expression quantification.

In addition, HiFi-slide results suggest a subtle downregulation in microglia expressions, which may be a result of the displacement of microglia cells during the neuron degeneration, and the spatial spreading patterns of the cell migration.

### Cell-Type Proximity Analysis

To further demonstrate the spatial analysis performed with HiFi-slides. The spatial proximity of cell types is compared between ND and AD samples. We used log2FC between the observed and expected cell type proximity for each cell type pair of ND and AD samples. Enrichment values were plotted and compared (Figure 5), where the higher value indicates the higher proximity (closer) in the AD spatial pattern. In order to systematically compare the two samples, we compare the enrichment values across cell type pairs, which is the log2FC between the observed and expected cell type proximities for each cell type pair. Being a log-scaled value, we extracted a log2FC of the enrichment as “enriched AD -enriched control” for each cell type pair. In Figure 5, we report the enrichment of the log2FC values.

**Fig 5.**
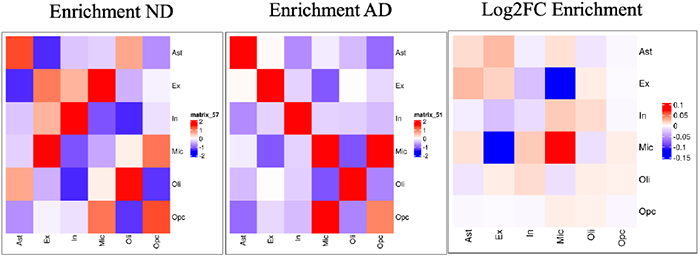
Spatial proximity analysis of AD vs. ND gene expression enrichment. Spatial proximity analysis was done with Giotto (35), with the log2FC enrichment values shown as a heatmap of the logarithm value.

A notable increase is observed in the spatial proximity among microglia cells for the AD sample, which suggests the morphology change and the ramification to amyloids. There is also an increase in the spatial proximity between astrocytes and excitatory neurons, showing the excitotoxicity activity increasing the spatial gathering of astrocytes to the neurons (33), and the astrocytes replacing the degenerated excitatory neurons, increasing the spatial proximity.

A noticeable decrease is observed in the spatial proximity between excitatory neurons and microglia cells in the AD case, as well as the increase between inhibitory neurons and microglia cells. The observation may implicate the migration of Mic groups in response to amyloid plaques. At the same time, some of the inhibitory neurons have higher resistance to the plaque (34), therefore showing higher proximity to the microglia cell groups.

## Methods

Formulas for reaction mixes are attached in Sup. Note 2. Sequences and oligonucleotide designs are attached in Sup. Note 3. Commercially available materials and consumables are attached in Sup. Note 4. The modularized protocols can be located on the protocols.io website (36).

### A. Slide Selection and Preparation

To be recycled with the HiFi-slides application, the used flow cell must be provided with the original sequencing files and a paired-end read configuration. The flow cells were screened based on the cluster density to avoid over or under-clustering strands. In addition, read 1 cannot involve custom read primer and must be over 70 bp. Selected Illumina NextSeq P3 flow cells were stored at 4 °C.

Dremel handheld Rotary tools with a diamond glass cutter were used to cut the adhesive regions of the flow cell to isolate the transparent surfaces. The surfaces were collected and placed in chambered slides for storage. Two surfaces were treated in a 2 mL tube with 1.5 mL 500 nM P7 primer solution prepared in Low TE with a 400rpm vortex to improve surface yield. The slides were immersed into the solution and incubated at 95 °C for 5 minutes. The tube was cooled at room temperature for 10 minutes. The surfaces were collected and placed in chambered slides. The slides were centrifuged to dry in a 1.5 mL microcentrifuge tube.

The slides were placed into the tube with prechilled BbSI Mix. They were incubated overnight at 37 °C and gently vortex (400rpm). On collection, the slide was placed on a chambered slide and washed with pipetting water on the surfaces three times. The slides were then centrifuged to dry, immersed in a 1.5 mL T4 ligation mix, and chilled at room temperature. The tube was incubated at 16 °C overnight. The slides were centrifuged to dry and placed in a cell dish for storage.

### B. Tissue Application

Fresh frozen human brain cortex samples were stored at -80 °C before spatial application. The samples were immersed in an OCT medium with a mold. The samples were then sectioned into 10-um slices and mounted on HiFi-slides with a cryostat. The slide was immersed in pre-chilled methanol for 20 minutes in a 1.5 mL tube. The slide was then retrieved from the tube and allowed to air dry. For the H & E staining, pipetting was done from the corner of the slides to avoid damaging the tissues.

The staining procedure was accomplished by pipetting just enough volume of the staining reagent to cover the sample and washing it with water each time after incubation. The slide was incubated with Hematoxylin for 10 minutes, then with a Bluing buffer for 7 minutes, and finally with Eosin for 2 minutes. The slices were scanned under a bright field with Keyence scanning microscope.

The tissue digestion mix was prepared, and the slides were transferred into chambered slides. The chambers were sealed, and the slides were incubated with 100 uL digestion mix for each slide at 37 °C for 12 minutes. The reaction mixture was then removed, and the slide surface was washed with 0.1X SSC solution before the reverse transcription was done by adding the RT mix and incubating overnight at 42 °C with a tilted ProFlex PCR system.

The reaction mix was removed on the second day, and the slide surface was washed with Ultrapure water. Exonuclease I treatment was performed by adding the Exon Mix and incubating for 3 hours. The reaction mix was removed using a pipette, and the slides were placed in a cell dish for temporary storage. The slides were washed with 0.1 N NaOH solution three times to remove any unwanted substances. The solution was then removed, and the slides were washed three times with 200 mM Tris-HCl solution, followed by three washes with water. The slides were then centrifuged to dry.

The Second-strand mix was then added to the slides on a tilted ProFlex PCR system and incubated at 37 °C for 3 hours. After incubation, the solutions were removed, and the slides were washed with water three times and then centrifuged to dry. The slides were eluted with 50 uL 0.1 N NaOH solution twice with 10 minutes of room temperature incubation for each round. Each slide’s total collection of 100uL elution was neutralized with 10 uL 3 M Sodium Acetate and purified with the NEB Monarch columns, with 50 uL of the purified single-stand library.

The slides were then moved to a 1.5 mL tube and were washed with 200 mM Tris-HCl solution three times, followed by three washes with water. The slide second strand synthesis was repeated four times with different barcodes conjugated with the second strand random primer. This step resulted in 4 50 uL single-strand libraries for each slide.

### C. Library Processing

The primer mix was combined with eluted DNA and NEB Q5 Master Mix to prepare the PCR reaction (50 uL DNA input, 75 uL Q5 Mater Mix, and 25 uL primer mix for each slide). The reaction was performed. The PCR products were purified with Monarch PCR Cleanup Column, with 80uL elution per second strand product.

The products were then quantified on Qubit. The library went through a size selection from 500 to 1500 bp by eluting from gel electrophoresis and quality control with TapeStation high sensitivity dsDNA assay. The libraries were sequenced with NovaSeq SP at 100 bp for Read 1 and 200 bp for Read 2 configuration.

### D. Data Processing

The data processing pipeline is composed of three main parts: 1) processing of the spatial barcodes (R1 of the recycled flow cell library), 2) processing of the HiFi-Slide read pairs (HiFi-Slide library: R1 is the spatial end, R2 is the RNA end), and 3) selection of the Region of Interest (ROI).

#### 1. Processing of the spatial barcodes

First, spatial barcodes are deduplicated. At this step, unique reads by sequence are selected and stored in a fasta file, and bwa indexes are created from their sequences. In addition, read names are labeled with a numerical identifier at the end indicating how many duplicates each read has (i.e., reads with identical sequences but different spatial locations). When duplicates are present, they are stored in a two-column tab-separated file, with the reference “unique” read in the first column and the duplicates in the second column, with as many rows per each unique read as its number of duplicates.

#### 2. Processing of the HiFi-Slide read pairs

HiFi-Slide R2 ends (RNA ends) are first preprocessed to filter out R2 ends overlapping R1 ends with more than 10 bp (default, PEAR v0.9.6) (37), too short reads, and then to trim Illumina adapters, polyG and polyX tails (fastp v0.23.2) (38). After these steps, R2 ends are aligned towards the human hg38 reference genome using STAR (v2.5.4b) (39), with the annotation file gen-code.v41 (–sjdbGTFfile) (40), and allowing all the alignments as output (–outFilterScoreMinOverLread and – outFilterMatchNminOverLread equal 0). Uniquely mapped reads are then selected (samtools v1.8) (41) and eventually annotated with the corresponding overlapping genes (one-base overlap).

Next, HiFi-Slide R1 ends (spatial ends) are aligned towards the spatial barcodes using bwa (v0.7.17)(42), and then filtered to remove the aligned R1 reads whose corresponding R2 ends were not mapped over the genome (previous step), to reduce both computational workload and memory occupation. Furthermore, we collect R1 reads mapped to spatial barcodes with the highest alignment score (AS). If an R1 read is aligned to *m* multiple barcodes that are tied at the highest score, all of these *m* alignments are considered.

In addition, each of these *m* barcodes could be found at *n* multiple locations (spots) over the flow cell (see deduplication in point 1 above). Thus, each R1 read could be potentially located at 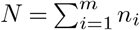 spatial locations. We filter out R1 reads with *N* >= 1000. Next, leveraging the HiFi-Slide read ID, the outputs from HiFi-Slide R1 and R2 processing are merged to obtain a tab-separated file with both spatial and gene information for each HiFi-Slide read pair. This file contains unique “read-gene” pairs in each line.

In the last step, we go from “read-gene” pairs to a final output tab-separated file where each row contains a unique “spot-gene” pair, with the gene expression level calculated as a “weighted read count,” which considers the potentially multiple locations of the read pairs. For read pairs aligned to barcodes with *N* > 1, we scale their gene expression level by *N*.

#### 3. ROI selection

Data within tiles of a predefined Region of Interest (ROI) are selected and will go under downstream analysis. Tiles under ROI are visually chosen from the tissue image. This step is necessary to select data coming from the tissue region (the tissue does not cover the entire flow cell, meaning some of the spatial locations are not actually under the tissue area) and to reduce computational workload.

### E. Data Analysis

#### 1. Cell type assignment to spots

To assign cell types to spots, we leveraged a set of 197 marker genes downloaded from the Human Protein Atlas (43), single cell type section, category “cell_type_enriched” (at least a four-fold higher mRNA level in a certain cell type compared to any other cell type), for the following cell types: astrocytes (Ast, 35 markers), excitatory neurons (Ex, 23 markers), inhibitory neurons (In, 25 markers), microglia (Mic, 12 markers), oligodendrocytes (Oli, 80 markers), oligodendrocyte precursor cells (Opc, 22 markers).

If a spot expresses one or more markers of a cell type, it is assigned to that specific cell type. If a spot expresses marker genes associated with several cell types, it is assigned to the cell type corresponding to the marker with the highest expression level. If a spot expresses marker genes associated with several cell types and two or more marker genes have the same highest expression level, then it is not assigned.

#### 2. Spatial clustering

The software Spaceflow (44) is used to perform spatial clustering. We ran Spaceflow on all the spots assigned to cell types (centroids) using the 197 cell type marker genes that are expressed using default parameters. To test for the association between spatial domains and cell types across spots, we performed a chi-square test for each “cell type-domain” pair using a contingency table where rows are “in the spatial domain” and “not in the spatial domain,” and columns are “in cell type” and “not in cell type.” In each entry is the number of spots.

Next, we generated a “cell type-by-domain” matrix with the odds ratios and the corresponding p-values, then we selected those significant entries whose odds ratio > 1 and p-value < 0.01. This allows us to associate each cluster with a group of cell types statistically.

#### 3. Cell type spatial proximity

The software Giotto (35) was used to evaluate the spatial proximity enrichment or depletion between pairs of cell types. The tool takes as input the gene-spot expression matrix, together with the cell type annotations and spatial coordinates of the spots, and it creates a spatial network connecting spots based on their physical distance allowing the evaluation of the spatial proximity enrichment or depletion between pairs of cell types.

## ACKNOWLEDGEMENTS

Research reported in this publication was supported by The Human BioMolecular Atlas Program (HuBMAP) of the National Institutes of Health under award number UH3CA256960. The content is solely the responsibility of the authors and does not necessarily represent the official views of the National Institutes of Health.

Recycled flow cells were provided by Ren Lab, Department of Cellular and Molecular Medicine, University of California San Diego School of Medicine, La Jolla, CA, USA, and Department of Cellular and Molecular Medicine, University of California San Diego School of Medicine, La Jolla, CA, USA.

Human cortex samples were provided by Banner Health Arizona.

This publication includes data generated at the UC San Diego IGM Genomics Center utilizing an Illumina NovaSeq 6000 that was purchased with funding from a National Institutes of Health SIG grant (#S10 OD026929).

Demonstration figures created with BioRender.com.

---

**Supplementary Table 1:**
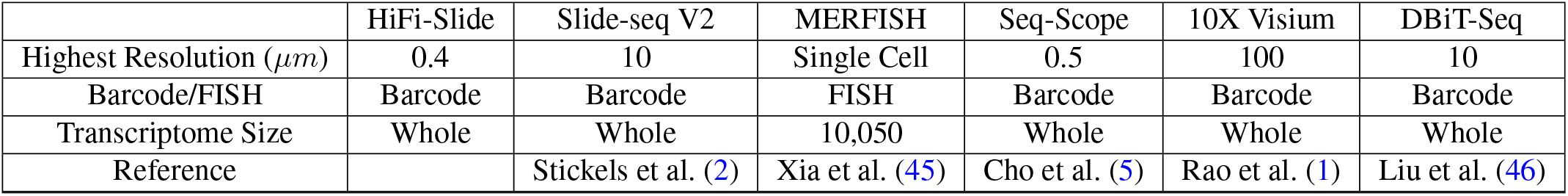
Transcriptomic State-of-Art Comparison

|  | HiFi-Slide | Slide-seq V2 | MERFISH | Seq-Scope | 10X Visium | DBiT-Seq |
| --- | --- | --- | --- | --- | --- | --- |
| Highest Resolution ($\mu m$) | 0.4 | 10 | Single Cell | 0.5 | 100 | 10 |
| Barcode/FISH | Barcode | Barcode | FISH | Barcode | Barcode | Barcode |
| Transcriptome Size | Whole | Whole | 10,050 | Whole | Whole | Whole |
| Reference |  | Stickels et al. (2) | Xia et al. (45) | Cho et al. (5) | Rao et al. (1) | Liu et al. (46) |

**Supplementary Table 2:** Reaction Mixes Preparation

|  |  |
| --- | --- |
| | BbSI Mix (50μl per reaction) |
| BbSI (NEB, R0539) | 1μl |
| 10X NEB Buffer r2.1 | 5μl |
| | T4 Ligation Mix (20μl per reaction) |
| T4 DNA Ligase Buffer (10X) | 2μl |
| T4 DNA Ligase (NEB, M0202) | 1μl |
| dT Solution (5μM) | 1μl |
| Water | 16μl |
| | Digestion Mix (10μl per reaction) |
| Pepsin reconstituted in 0.1M HCl | 5μl |
| Water | 5μl |
| | RT Mix (20μl per reaction) |
| Maxima H Minus Enzyme Mix (TF, K1652) | 1μl |
| Maxima H Minus Buffer (RT Buffer) | 4μl |
| 10mM dNTP | 1μl |
| Water | 14μl |
| (Optional) Visual TSO 10μM Oligo | 1μl, diluted in water |
| | Exon Mix (20μl per reaction) |
| Exonuclease I (E.coli)(NEB, M0293) | 2μl |
| Exonuclease I Reaction Buffer | 2μl |
| Water | 16μl |
| | Second Strand Mix (20μl per reaction) |
| Kelnow Frag, Exo- (NEB, M0212) | 2μl |
| NEBuffer 2 | 2μl |
| 10mM dNTP | 2μl |
| Random Primer (100μM) | 1μl |
| Water | 13μl |
| | Primer Mix (100μl per reaction) |
| P5 Primer Stock Solution (100μM) | 3μl |
| P7 Primer Stock Solution (100μM) | 3μl |
| Water | 94μl |
| | PCR Reaction Mix (50μl per reaction) |
| Primer Mix | 10μl |
| Eluted DNA | 15μl |
| Kapa HIFI Hotstart ReadyMix (Roche Diagnostics, KK2602) | 25μl |

**Supplementary Table 3:**
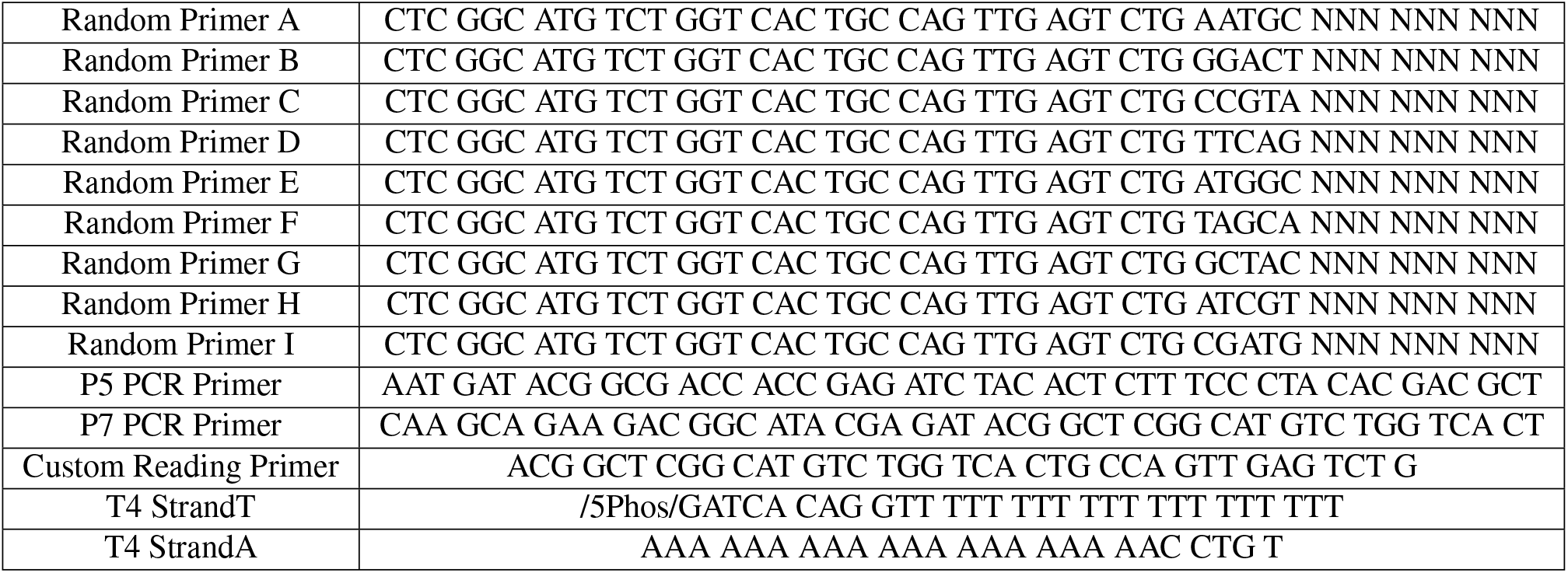
HiFi-Slide Oligo Sequences

**Supplementary Table 4:**
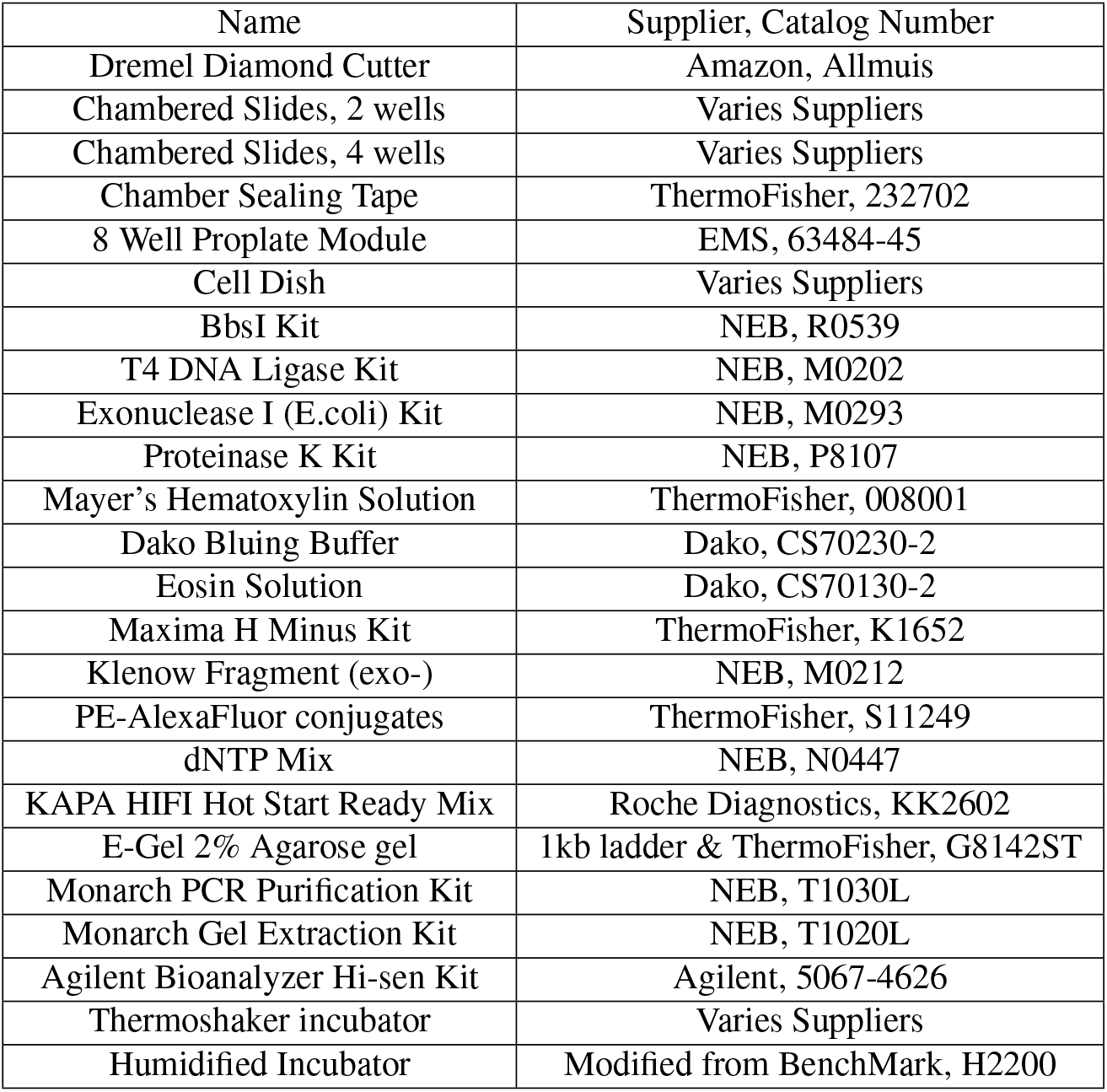
HiFi-Slide Material Checklist

| Name | Supplier, Catalog Number |
| --- | --- |
| Dremel Diamond Cutter | Amazon, Allmuis |
| Chambered Slides, 2 wells | Varies Suppliers |
| Chambered Slides, 4 wells | Varies Suppliers |
| Chamber Sealing Tape | ThermoFisher, 232702 |
| 8 Well Proplate Module | EMS, 63484-45 |
| Cell Dish | Varies Suppliers |
| BbsI Kit | NEB, R0539 |
| T4 DNA Ligase Kit | NEB, M0202 |
| Exonuclease I (E.coli) Kit | NEB, M0293 |
| Proteinase K Kit | NEB, P8107 |
| Mayer's Hematoxylin Solution | ThermoFisher, 008001 |
| Dako Bluing Buffer | Dako, CS70230-2 |
| Eosin Solution | Dako, CS70130-2 |
| Maxima H Minus Kit | ThermoFisher, K1652 |
| Klenow Fragment (exo-) | NEB, M0212 |
| PE-AlexaFluor conjugates | ThermoFisher, S11249 |
| dNTP Mix | NEB, N0447 |
| KAPA HIFI Hot Start Ready Mix | Roche Diagnostics, KK2602 |
| E-Gel 2% Agarose gel | 1kb ladder & ThermoFisher, G8142ST |
| Monarch PCR Purification Kit | NEB, T1030L |
| Monarch Gel Extraction Kit | NEB, T1020L |
| Agilent Bioanalyzer Hi-sen Kit | Agilent, 5067-4626 |
| Thermoshaker incubator | Varies Suppliers |
| Humidified Incubator | Modified from BenchMark, H2200 |

